# Divergent Energy Expenditure Impacts Mouse Metabolic Adaptation to Acute High-Fat/High-Sucrose Diet Producing Sexually Dimorphic Weight Gain Patterns

**DOI:** 10.1101/840702

**Authors:** E. Matthew Morris, Roberto D. Noland, Julie A. Allen, Colin S. McCoin, Qing Xia, Devin C. Koestler, Robin P. Shook, John R.B. Lighton, Julie A. Christianson, John P. Thyfault

## Abstract

**Objective:** Long-term weight gain can result from cumulative small weight increases due to short-term excess caloric intake during weekends and holidays. Increased physical activity may mediate weight gain through increases in energy expenditure (EE) and reductions in energy balance. Current methods for modulating mouse EE (e.g. – exercise, chemical uncouplers, etc.) have confounding effects. However, it is known that mouse EE linearly increases as housing temperature decreases below the thermoneutral zone.

**Methods:** To determine how robust differences in baseline EE impact 7-day changes in weight and body composition on low-fat and high-fat, high-sucrose (HFHS) diets, we performed indirect calorimetry measurements in male and female mice housed at divergent temperatures (20°C vs. 30°C).

**Results:** As expected, mice housed at 30°C have ∼40% lower total EE and energy intake compared to 20°C mice regardless of diet or sex. Energy balance was increased with HFHS in all groups, with ∼30% greater increases observed in 30°C versus 20°C mice. HFHS increased weight gain regardless of temperature or sex. Interestingly, no HFHS-induced weight gain differences were observed between females at different temperatures. In contrast, 30°C male mice on HFHS gained ∼50% more weight than 20°C males, and ∼80% more weight compared to 30°C females. HFHS increased fat mass across all groups but 2-fold higher gains occurred in 30°C mice compared to 20°C mice. Females gained ∼35% less fat mass than males at both temperatures.

**Conclusions:** Together, these data reveal an interaction between divergent ambient temperature-induced EE and sex that impacted diet-induced patterns of short-term weight gain and body composition.

**Highlights:**

- Utilized ambient temperature differences as an experimental tool to study the impact of divergent baseline energy expenditure on metabolic adaptation to high-fat, high-sucrose diet.
- Baseline energy expenditure and sex interact to impact diet-induced changes in body composition and weight gain.
- The energy expenditure and sex interaction is a result of an inverse relationship between fat mass gain and weight-adjusted total energy expenditure, as well as, diet-induced non-shivering thermogenesis.
- These data support that the hypothesis that higher energy expenditure amplifies the coupling of energy intake to energy expenditure during energy dense feeding, resulting in reduced positive energy balance and reduced gains in weight and adiposity.
- First evidence that energy expenditure level plays a role in the composition of weight gained by female mice during acute HFHS feeding.
- This study further highlights issues with obesity/energy metabolism research performed in mice at sub-thermoneutral housing temperatures, particularly with sex comparisons.

**GRAPHIC ABSTRACT:** **Legend:** Male and female mice housed at 30°C had lower energy expenditure (EE) & energy intake (EI), while having greater energy balance (EB), during 7-day high-fat/high-sucrose (HFHS) feeding compared to male and female mice, respectively, housed at 20°C. However, female mice had lower EB compared to males at both housing temperature. Female mice housed at 30°C gained less weight than 30°C males but gained the same relative amount of fat mass during acute HFHS feeding. Interestingly, 20°C females gained the same amount of weight as 20°C males but gained primarily fat-free mass, while the males gained the same proportion of fat as 30°C males and females.

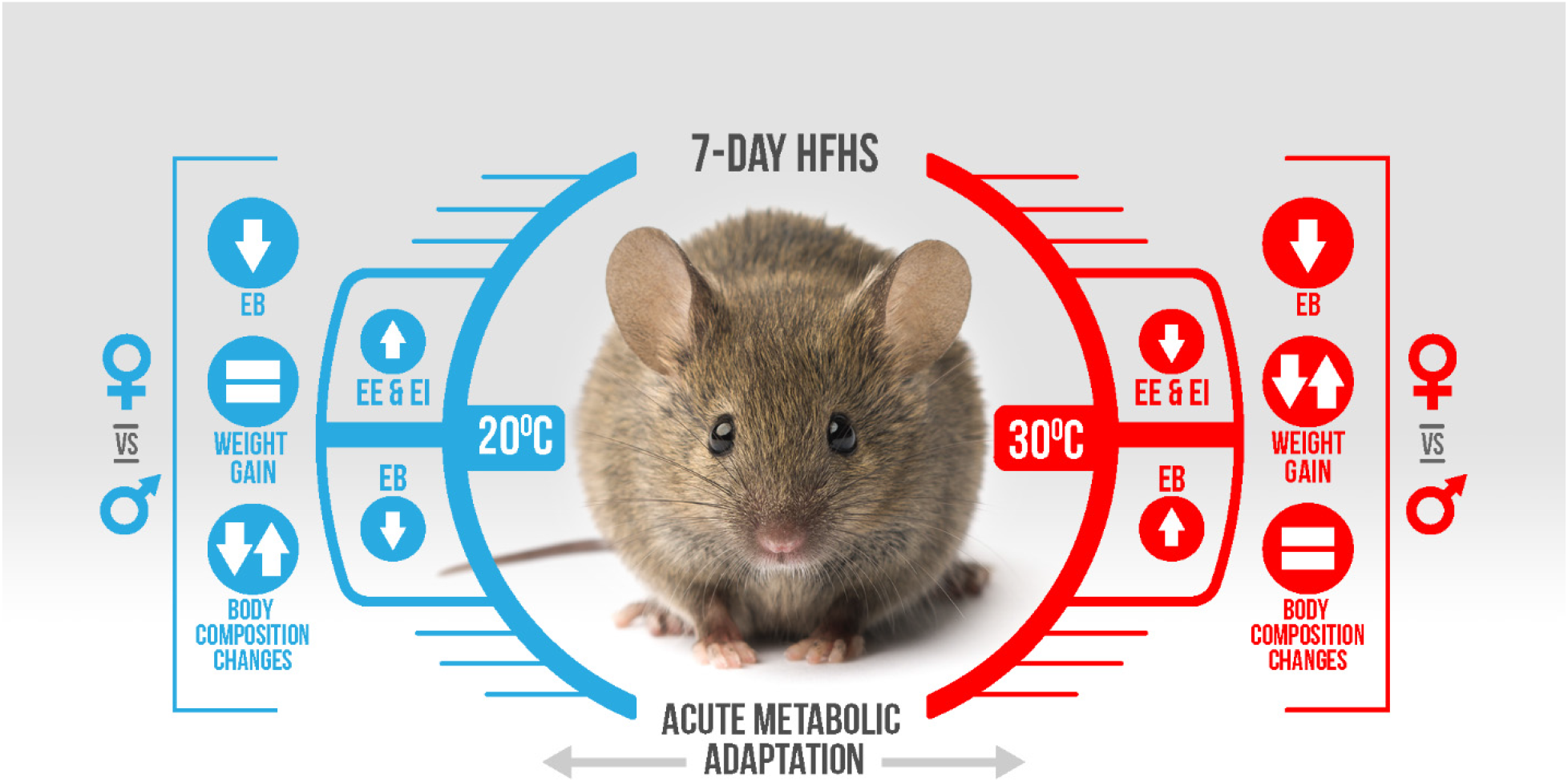

## 1. INTRODUCTION

Obesity can occur through episodic periods of weight gain caused by consumption of energy dense foods during weekends, holidays, or seasons [1–5]. Simply stated, weight gain occurs in a given time frame when the difference in energy intake exceeds energy expenditure resulting in a positive energy balance [6]. This positive energy balance represents a shift in the flux of energy consumed from expenditure to storage [6–8], comprised primarily of increased fat mass [9]. Previous research shows that a complex combination of hedonic, hormonal, metabolic, and sexually dimorphic regulatory mechanisms can alter energy intake and/or energy expenditure [10–13], but the integrated mechanisms that regulate episodic weight gain during acute periods of energy dense feeding remain unclear.

Current recommendations to prevent weight gain and treat obesity include increasing physical activity or daily exercise with a goal of increasing total energy expenditure (TEE) and improving energy balance [14–18]. Improved energy balance at higher physical activity levels is proposed to be achieved through greater coupled sensitivity of energy intake regulation to energy expenditure [6, 8]. However, increased physical activity produces multiple systemic and tissue-specific adaptations independent of EE (reviewed in [19–21]), complicating the direct investigation of modulating EE to protect against diet-induced weight gain. Additionally, observed sex differences in physical activity levels and physiological adaptation may impact weight gain prevention and/or obesity treatment [22, 23]. To more specifically study the impact of total EE (TEE) on acute diet-induced weight gain we leveraged the ability of divergent housing temperatures to cause grossly different EE in male and female mice before exposing them to a subsequent metabolic challenge (7-day high-fat/high-sucrose feeding). We utilized indirect calorimetry and EchoMRI to assess changes in energy metabolism, weight gain, and body composition to the diet. We hypothesized that higher TEE would provide protection against acute diet-induced weight gain, but that sexual dimorphic responses would emerge. We revealed that only male mice gained more weight at a low TEE, whereas females gained the same weight regardless of TEE. However, female mice at low TEE gained the same proportion of fat mass as males, while females at higher TEE gained primarily fat-free mass.

## 2. MATERIALS and METHODS

### 2.1 Animals

Male and female C57Bl/6J (#000664, Jackson Laboratories, Bar Harbor, ME, USA) mice (6-weeks old) were individually housed at either 20°C or 30°C on a reverse light cycle (light 10P – 10A) with *ad lib* access to low-fat, control diet [LFD, D12110704 (10% kcal fat, 3.5% kcal sucrose, 3.85 kcal/gm), Research Diets, Inc., New Brunswick, NJ, USA] for three weeks. At 9-weeks of age, animal weights and food weight was monitored prior to and following the 7 days of both LFD and the subsequent high-fat, high-sucrose diet [HFHS, D12451, 45% kcal fat, 17% kcal sucrose, 4.73 kcal/gm] at the assigned ambient temperature. At the end of the HFHS 7-day feeding, mice were food withdrawn for 2hrs (0800). The animal protocols were approved by the Institutional Animal Care and Use Committee at the University of Kansas Medical Center and the Subcommittee for Animal Safety at the Kansas City Veterans’ Hospital.

### 2.2 Body Composition Analysis

Body composition was measured by qMRI using the EchoMRI-1100 (EchoMRI, Houston, Texas, USA). Fat free mass (FFM) was calculated as the difference between body weight and fat mass (FM). Body composition was determined prior to, and after each of the 7-day feedings.

### 2.3 Indirect Calorimetry, Energy Metabolism, & Behavior Analysis

Starting at 9-weeks of age (n=12), energy metabolism was assessed at 20°C or 30°C ambient temperature for 7 days on LFD followed by 7 days of HFHS by measuring VO_2_ and VCO_2_ in a Promethion continuous metabolic monitoring system (Sable Systems International, Las Vegas, NV, USA), as described previously [24, 25]. Animals were acclimated to the indirect calorimetry cages for 5 days prior to initiation of data collection. Rate of energy expenditure was calculated with a modified Weir equation [EE (kcal/hr) = 60*(0.003941*VO2+0.001106*VCO2)], and respiratory quotient (RQ) as VCO_2_/VO_2_. Total energy expenditure (TEE) was calculated as the daily average rate of energy expenditure for each day times 24 and summed across the 7 days of each diet. Resting energy expenditure (REE) was calculated from the average rate of EE during the 30-minute period with the lowest daily EE as kcal/hr and extrapolated to 24hrs and summed across the 7 day dietary periods. Non-resting energy expenditure (NREE) was calculated as the difference between TEE and REE. 7-day average diurnal change in RQ was calculated as the difference in daily average of the dark cycle RQ minus the light cycle RQ for each diet. Diet-induced changes in RQ, REE, and NREE were calculated as the difference in the 7-day HFHS data minus the 7-day LFD data. Energy intake (EI) was calculated as the total food intake for each feeding period times the energy density of each diet. Energy balance (EB) was calculated as the difference between the total EI and TEE throughout each 7-day dietary exposure period. FI, EI, and EB data during HFHS feeding from two 20°C female mice was not included in data analysis due to excessive food spillage. Percent metabolic efficiency was calculated as: (change in fat mass (kcal) + change in lean mass (kcal)/EI, where the energy content for fat and lean mass was 9.32 kcal/g & 1.19 kcal/g, respectively [26]. Thermic effect of food (TEF) was determined from the consensus thermic effect of food for fat (2.5%), carbohydrate (7.5%), and protein (25%), and the manufacturer provided diet information for each diet [27, 28]. As such, the TEF for LFD (D12110704, Research Diets, 3.85 kcal/g, 10% kcals fat, 65% kcal carbohydrate, 20% kcals protein) is 10.5% or 0.4043 kcal/g, and HFHS (D12451, Research Diets, 4.73 kcal/g, 45% kcals fat, 35% kcal carbohydrate, 20% kcals protein) is 8.75% or 0.4139 kcal/g. This method of calculating TEF reduces the potential influence of neurobehavioral adaptations of the fed/fasted transition impacting changes in EE, through calculation of TEF across the entire 7-day period of each diet. Activity energy expenditure (AEE) was calculated as the difference between NREE and TEF. All_Meters is an assessment of cage activity including gross and fine movements; and is determined using the summed distances calculated from the Pythagoras’ theorem that the mouse moved since the previous data point based on XY second by second position coordinates. Cost of movement (CoM) was calculated as the AEE divided by total meters traveled over the 7-days of each diet. LFD data for 4 additional male and female mice at 30°C is included, no HFHS was collected for this subset as ambient temperature control was lost. All data from one 20°C female mouse was excluded after discovering malocclusion at necropsy.

### 2.4 Statistical Analysis

Data are presented as scatter plots with means and standard error. The two-standard deviation test was utilized to test for outliers in each group. Utilizing R statistical programming language version 3.5.1, (http://r-project.org), a series of linear mixed effects models were used to assess the relationship between anthropometric and energy metabolism measures with temperature (30°C/20°C), sex (Male/Female), and diet (LFD/HFHS). Linear effects models were fit to anthropometric and energy metabolism measures and included fixed effects terms representing the main effects of temperature, sex, and diet, all two-way interactions involving these terms and their three-way interactions, and a random intercept term for mouse to account for the anticipated autocorrelation given that multiple measurements were collect on the same mice. Models were additionally adjusted for fat mass and fat-free mass to control for their potential confounding effects. Adjusted means and partial eta-squared values as approximations of effect size were calculated. Additionally, parameter estimates obtained from the linear mixed models along with linear comparisons were conducted and adjusted for multiple comparisons using a Bonferroni correction. Main effects are discussed only when all pairwise treatment comparisons within that parameter were significant. Diet-induced changes were calculated as the difference in 7-day total of each variable during HFHS minus the LFD 7-day total. A two-way ANOVA was utilized to determine main effects of temperature and sex in diet-induced change data. Where significant main effects were observed, post hoc analysis was performed using least significant difference to test for any specific pairwise differences using SPSS version 25 (SPSS Inc., Armonk, New York). Statistical significance was set at p<0.05.

## 3. RESULTS

### 3.1 Systemic Energy Metabolism

Indirect calorimetry was utilized to investigate the role of differences in EE on systemic energy metabolism and HFHS-induced weight-gain in mice housed at 30°C versus 20°C. TEE was ∼40% lower in 30°C mice of both sexes and diets compared to 20°C (Figure 1A, p<0.0001), driving a significant 3-way interaction (p<0.0001). Additionally, HFHS feeding increased TEE ∼5-15% in all groups (p<0.0001). Female mice had lower TEE compared to males (p<0.05) except in the 20°C, HFHS-fed condition. 7-day EI was greater in 20°C mice on LFD and HFHS (∼55% & ∼32%, respectively) compared to 30°C regardless of sex (Figure 1B, p<0.0001). As expected, EI increased on HFHS compared to LFD across all groups (p<0.0001). The 30°C females and males increased EI 44% & 63%, respectively during HFHS feeding; while 20°C mice increased ∼34%. Females were observed to have lower EI (p<0.006) in all contrasts except 30°C LFD. EB was calculated as the difference in 7-day EI and TEE. While no difference in EB was observed during LFD feeding in either sex or temperature (Figure 1C), a significant increase in EB was induced by HFHS in all groups (p<0.001). Additionally, HFHS feeding resulted in lower EB in female mice housed at both 20°C & 30°C (∼30%, p<0.01). Further, 30°C housing produced greater EB during HFHS feeding in both male and female mice (25% & 45%, respectively, p<0.01).

**Figure 1:**
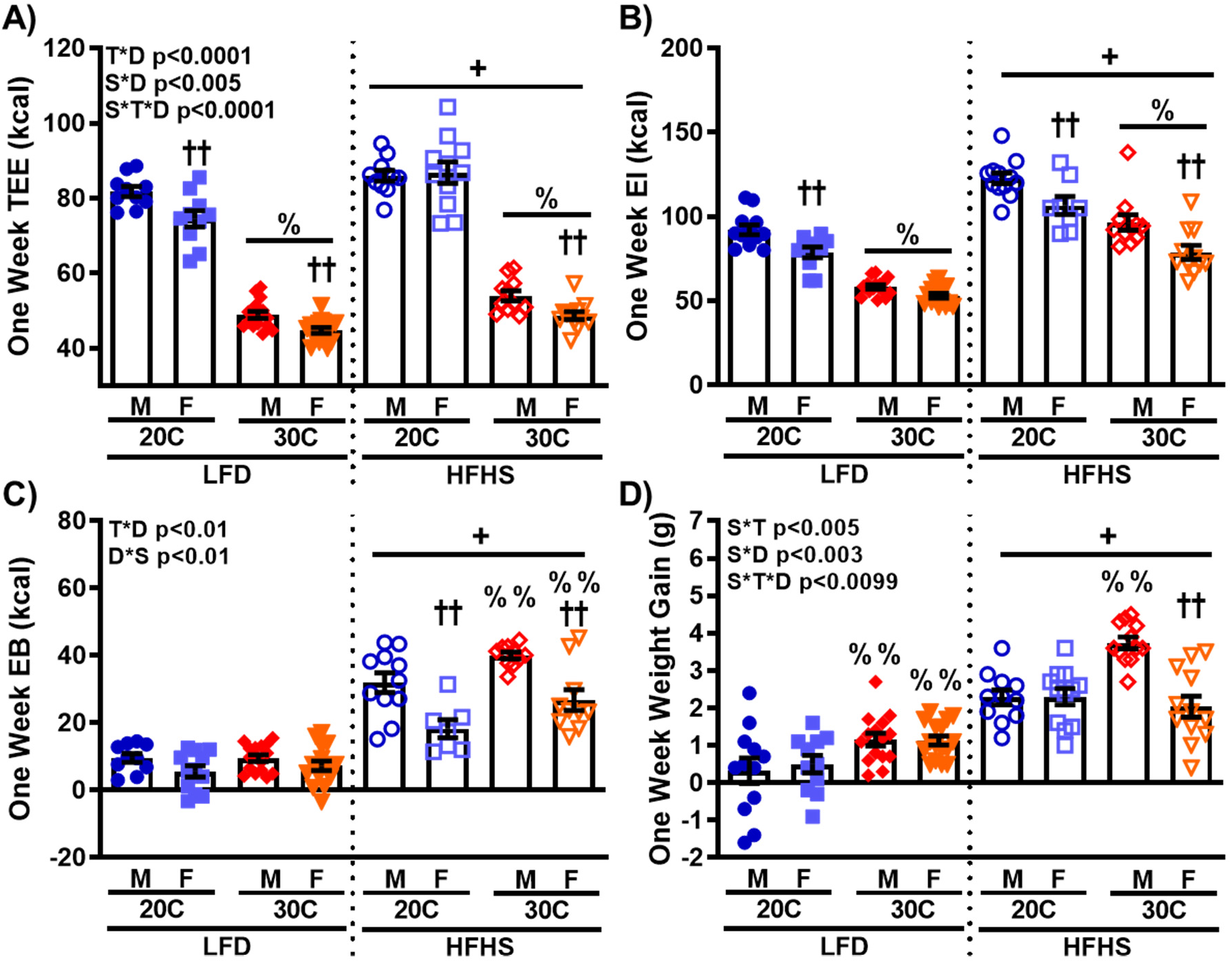
Divergent energy metabolism in mice due to different ambient housing temperature produces sexually dimorphic HFHS-induced weight gain. A) Indirect calorimetry was utilized to determine one week total energy expenditure (TEE) in male and female C57Bl/6J mice during 7-days of LFD followed by 7-days of HFHS (n=10-16). B) Energy intake (EI) during each dietary exposure was determined as the sum of food intake (g) times the energy density (kcal/g) for each diet (n=7-16). C) Energy balance was calculated as difference in EI and TEE for each mouse during each diet exposure (n=7-16). D) One week change in body weight (n=10-16). Values are means ± SEM. % p<0.05 main effect of 20°C vs. 30°C, + p<0.05 main effect of LFD vs. HFHS, †† p<0.05 male vs. female within temperature by diet group, %% p<0.05 20°C vs. 30°C within sex by diet group.

Initial body weights were not different between mice housed at 20°C vs. 30°C, with females weighing ∼25% less than males in both temperature groups. All initial, end of LFD (Day 7), and end of HFHS (Day 14) anthropometric data is presented in Supplemental Table A.1. Body weight gain during the 7-days of LFD was 3.6- and 2.3-fold higher (males and females, respectively) housed at 30°C compared to 20°C (Figure 1D, p<0.04). No difference in weight gain was observed between sexes on LFD at either temperature. Subsequent HFHS feeding resulted in a significant interaction of temperature by sex by diet (p<0.001). This interaction was primarily driven by the main effect of HFHS on weight gain regardless of temperature or sex (p<0.05). Interestingly, temperature did not effect weight gain in HFHS-fed female mice; whereas, HFHS-fed male mice housed at 30°C gained ∼65% more weight compared to 20°C males (p<0.05). Moreover, the 30°C male mice gained ∼80% more weight compared to females at 30°C (p<0.05).

### 3.2 Changes in Body Composition

We utilized qMRI to assess how baseline differences in EE impacted changes in body composition. Greater fat mass (FM) gain was observed at 30°C during the LFD (Figure 2A, p<0.0001) and a further increase was induced by HFHS feeding (p<0.0001). Transition to HFHS resulted in ∼50% & ∼70% less FM gain in female mice compared to male mice at 20°C & 30°C, respectively (p<0.0001). Importantly, 30°C housing resulted in much greater increases in FM on HFHS (64% & 2.8-fold increases in males and females, respectively) compared to 20°C (p<0.0001). In contrast to FM, temperature had no impact on changes in fat-free mass (FFM) during LFD (Figure 2B). Interestingly, FFM increased in female 20°C mice on HFHS compared to male 20°C (2.6-fold, p<0.0001) and female 20°C mice fed LFD (85%, p<0.005). Additionally, change in FFM was less in 30°C female mice compared to 20°C female and 30°C male mice (∼65%, p<0.0006 & ∼40%, p<0.04, respectively). To further highlight the interaction of sex, temperature, and diet in short-term weight gain and type of weight gained, Figure 2C displays the one week change in body weight as the components of FM and FFM gained. The metabolic efficiency was calculated as the sum of the stored energy from change in FM and lean mass divided by EI (Figure 2D) [26]. Calculated percent metabolic efficiency shows a similar pattern to the FM data (Figure 2A). Importantly, the difference in metabolic efficiency between temperature groups on HFHS increased to 2.2-fold for males & 3.7-fold for female mice (p<0.0001). Together, these data demonstrate an interaction of sex, temperature, and dietary exposure to impact changes in body composition, particularly, in female animals.

**Figure 2:**
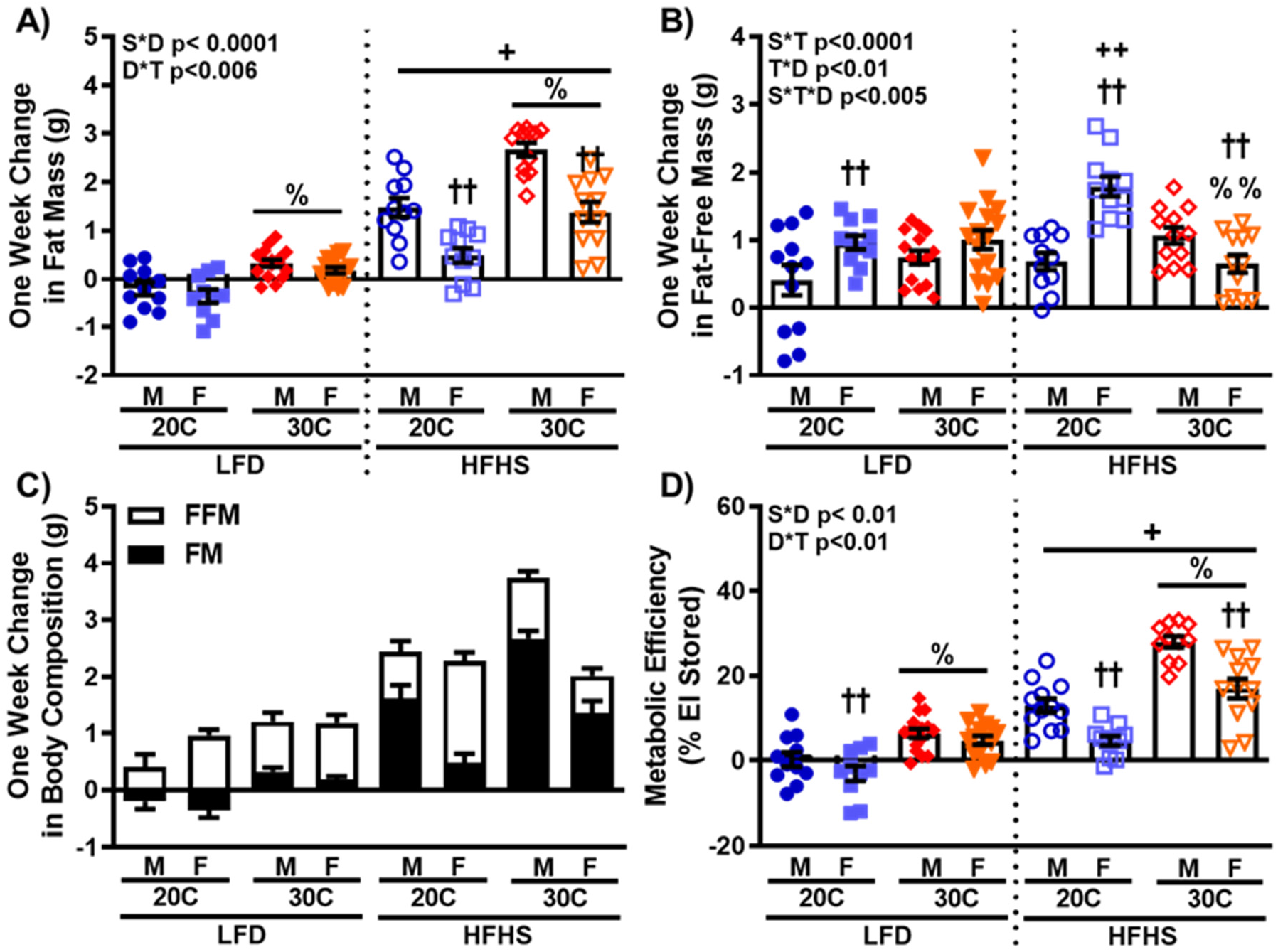
Higher energy metabolism during 20°C housing changes type of weight gained in female mice during one week HFHS feeding. Body composition analysis utilizing qMRI was performed before and after each diet exposure. The difference in the initial and final values during the one week of LFD and HFHS are displayed as (A) change in fat mass (FM) and (B) fat-free mass (FFM). C) One week change in body weight presented as change in FM and FFM. D) Metabolic efficiency as the percent of EI stored as FM and FFM. Values are means ± SEM. n=10-16. % p<0.05 main effect of 20°C vs. 30°C, + p<0.05 main effect of LFD vs. HFHS, †† p<0.05 male vs. female within temperature by diet group, %% p<0.05 20°C vs. 30°C within sex by diet group. ++ p<0.05 LFD vs. HFHS within temperature by sex group.

### 3.3 Component Analysis of TEE

To better characterize the interaction of temperature, sex, and diet on systemic energy metabolism, TEE was dissected into resting (REE) and non-resting (NREE) components (Figure 3A & B, respectively). Where REE is primarily comprised of basal metabolic rate and non-shivering adaptive thermogenesis, and NREE encompasses the thermic effect of food and activity-induced EE. A significant 3-way interaction of temperature*sex*diet (p<0.0009) was observed for REE, driven by the ∼55% reduction in REE in 30°C mice regardless of sex or diet (p<0.0001). HFHS feeding resulted in an ∼15-25% increase in REE across all groups (p<0.0001). Female mice had ∼10% lower NREE compared to males on LFD (p<0.02). The transition to HFHS reduced NREE ∼15% in 20°C male and female mice (p<0.0001). However, 30°C mice did not lower NREE after a transition to the HFHS. TEE is graphically represented in Figure 3C as the components REE and NREE to more clearly visualize each component’s absolute amount during different temperature and diet conditions. Additionally, the percent of TEE for REE and NREE is presented in Figure 3D. Main effects of temperature (p<0.0001) and diet (p<0.0001) were observed for percent REE regardless of sex. For both sexes, REE comprised ∼70% of TEE at 20°C compared to ∼53% at 30°C on LFD. On HFHS, REE comprised ∼78% and 58% of TEE for 20°C and 30°C, respectively.

**Figure 3:**
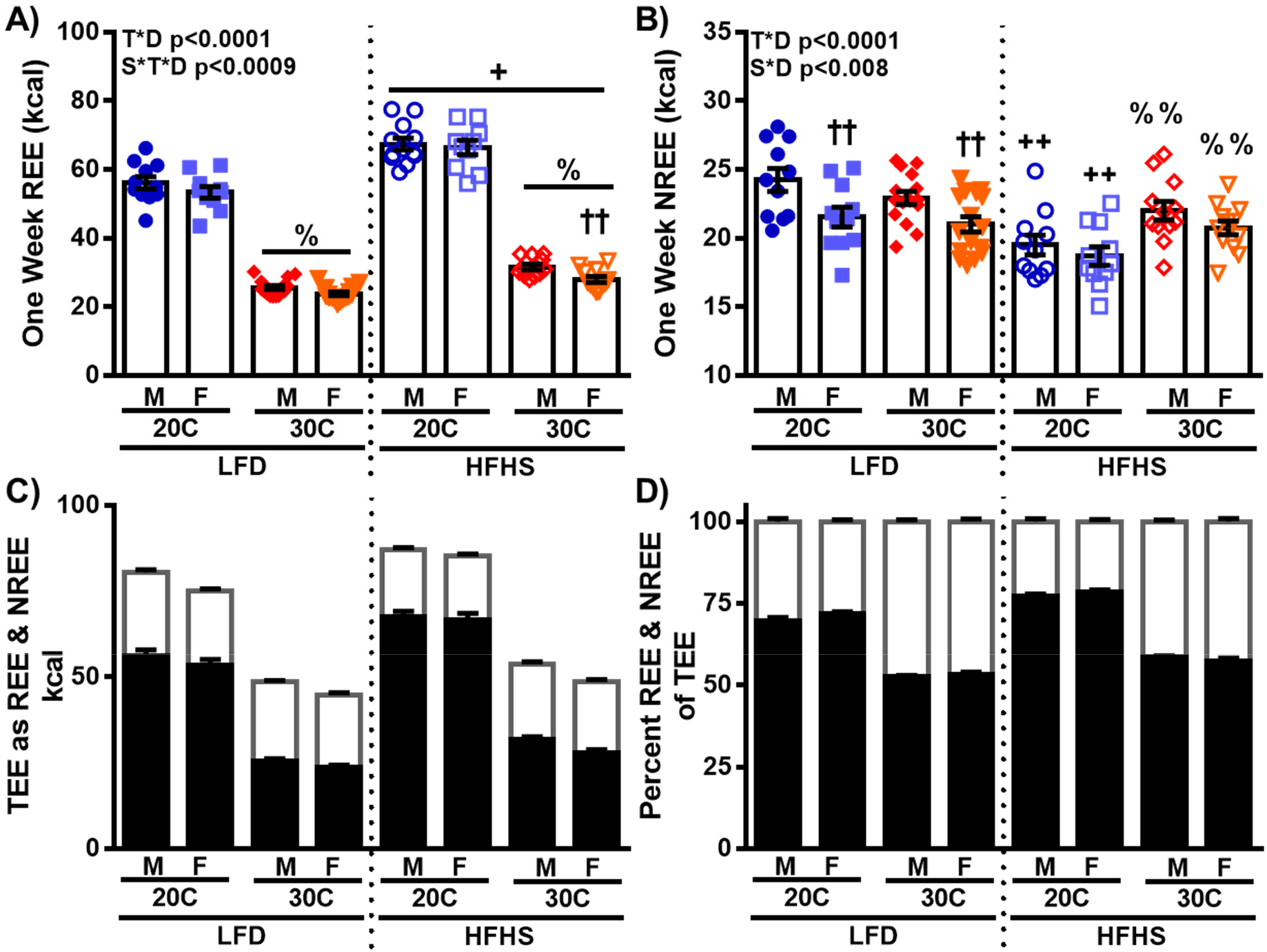
Component analysis of total energy expenditure. A) Indirect calorimetry was utilized to determine one week resting energy expenditure (REE) in male and female C57Bl/6J mice during 7-days of LFD followed by 7-days of HFHS. B) Non-resting energy expenditure (NREE) was calculated as the difference between TEE and REE over the 7-day dietary exposures. C) TEE represented as the components: ■ REE & □ NREE. D) Percent of TEE comprised of ■ REE & □ NREE. Values are means ± SEM. n=10-16. % p<0.05 main effect of 20°C vs. 30°C, + p<0.05 main effect of LFD vs. HFHS, †† p<0.05 male vs. female within temperature by diet group, %% p<0.05 20°C vs. 30°C within sex by diet group, ++ p<0.05 LFD vs. HFHS within temperature by sex group.

### 3.4 Co-variate Analysis of Energy Metabolism

Co-variate analysis of the energy metabolism outcomes was performed to assess the effect of differences in the components of body weight (FM and FFM) on the interpretation of TEE and EI. Adjusted estimated marginal means and partial eta squared values are shown in Figure 4. Following ANCOVA to adjust for differences in fat- and fat-free mass, females had higher TEE in all diet X temperature comparisons (Figure 4A, p<0.01). Main effects of temperature and diet were observed as in the absolute TEE data (Figure 1A). Importantly, FM was not a significant co-variate of TEE; while FFM was significant (p<0.001) and showed a moderate effect size (partial eta-squared – 0.38) on TEE. Adjustment of EI removed all differences between males and females (Figure 4B), while maintaining the previously observed (Figure 1B) main effects for temperature and diet. Again, FM was not a significant co-variate, and while FFM was significant (p<0.05), a very small effect size was observed (partial eta-squared – 0.07).

**Figure 4:**
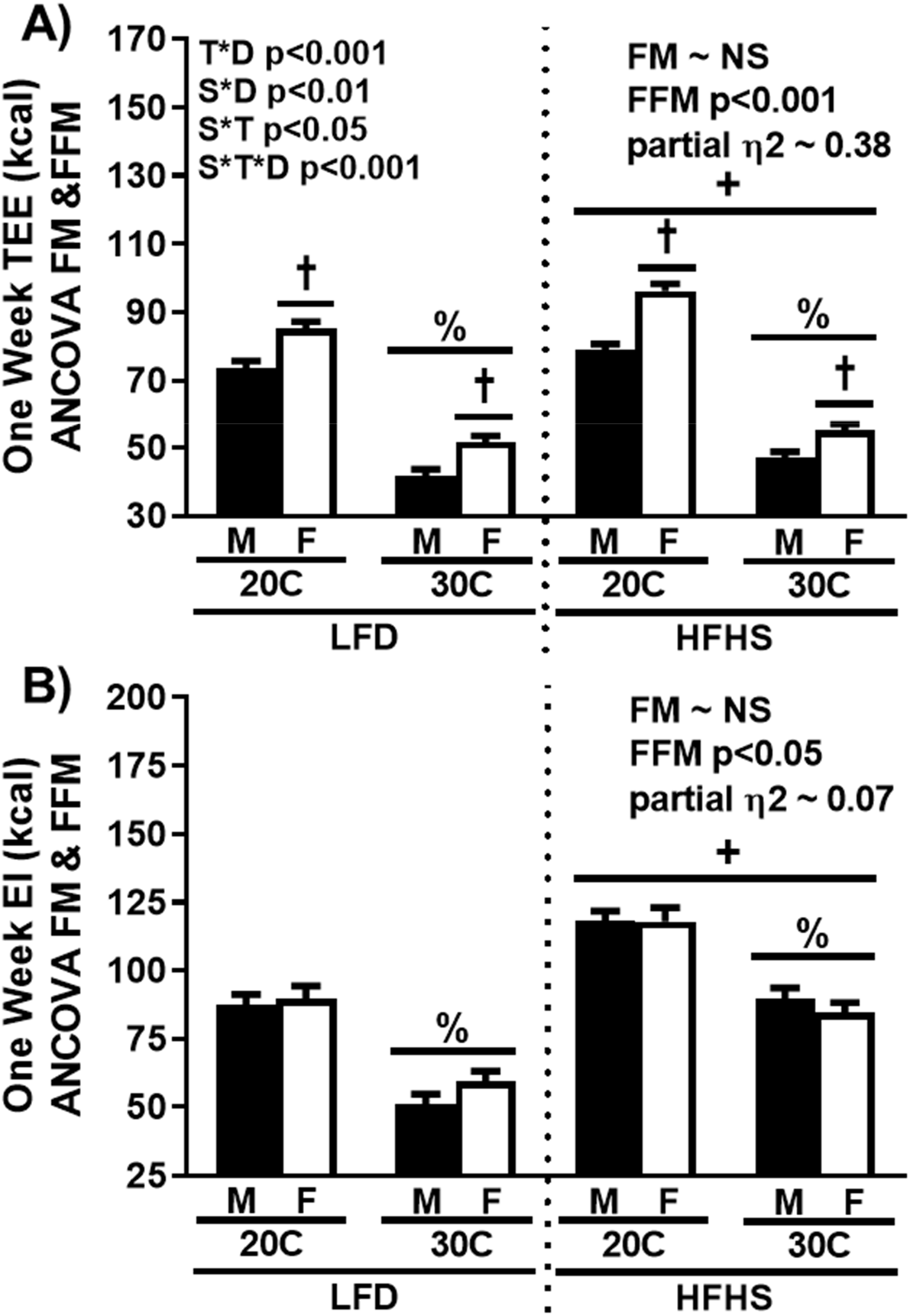
Female mice have greater total energy expenditure and equal energy intake following co-variate analysis by fat and fat-free mass. A) TEE (n=10-12) and B) EI (n=8-12) following ANCOVA for fat mass + fat-free mass is expressed as the estimated mean ± SEM. Estimated effect size of the significant covariates for each ANCOVA analysis are presented as partial eta squared. % p<0.05 main effect of 20°C vs. 30°C, + p<0.05 main effect of LFD vs. HFHS, † p<0.05 main effect of male vs. female.

### 3.5 HFHS-induced Changes in Energy Metabolism

To further assess the roles of temperature and sex on diet-induced changes in energy metabolism, we quantified substrate utilization (respiratory quotient, RQ), metabolic flexibility, and within animal EE adaptations following the transition to HFHS. Daily average respiratory quotient (RQ) was significantly reduced in all groups fed HFHS as expected (Figure 5A, p<0.0001). No significant contrasts were observed for temperature during either LFD or HFHS feeding, and only 30°C females were significantly lowered by HFHS compared to male mice (p<0.004). We calculated two measures of metabolic flexibility, which represents the capacity to adapt substrate utilization based on changes in physiological state [10, 29]. First, we quantified the daily average difference between dark and light cycle RQ (Figure 5B). LFD mice at 30°C have reduced change in diurnal RQ compared to 20°C (p<0.005); demonstrating that mice housed at 30°C are inherently less metabolically flexible. Additionally, HFHS further lowered average diurnal RQ (p<0.002) in all groups; indicating that short-term HFHS feeding was sufficient to exacerbate metabolic inflexibility. Interestingly, no difference in sex was observed across any of the comparisons. Second, the capacity of diet to alter substrate utilization was also assessed as the change in daily average RQ from LFD to HFHS (Figure 5C). 30°C mice showed a smaller HFHS diet-induced reduction in daily average RQ compared to 20°C mice (p<0.0001). However, 30°C female mice had greater HFHS diet-induced changes in RQ compared to males (p<0.02). Figure 5D shows the average change in daily RQ during the transition from LFD to HFHS. The figure highlights the rapid RQ decrease in all groups and slower transient response of the 30°C mice.

**Figure 5:**
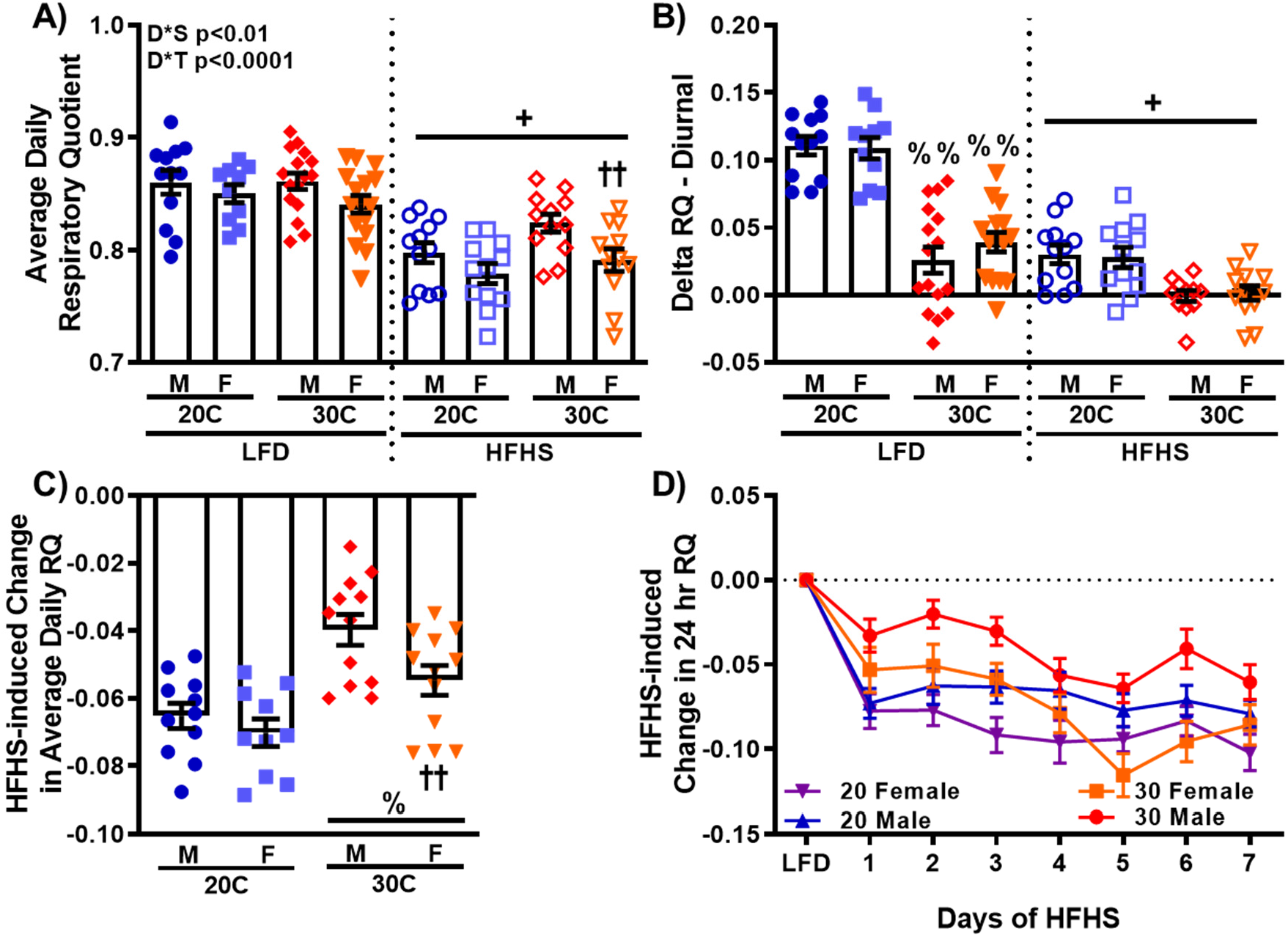
Divergent energy metabolism in mice due to different ambient housing temperature produces differences in metabolic flexibility on both low-fat and high-fat/high-sucrose diets. A) Indirect calorimetry was utilized to determine average daily respiratory quotient (RQ). Metabolic flexibility was assessed as: B) the daily difference between dark cycle RQ minus light cycle RQ averaged across the dietary exposures and C) the difference in average daily RQ during HFHS feeding and average daily RQ during LFD feeding. D) RQ change from LFD RQ during each day of the HFHS exposure. Values are means ± SEM. n=10-16. + p<0.05 main effect of LFD vs. HFHS, % p<0.05 main effect of 20°C vs. 30°C, †† p<0.05 male vs. female within temperature by diet group.

Diet-induced non-shivering thermogenesis is the adaptive capacity to increase EE in response to increases in EI and is a compensatory mechanism for limiting increased EB during transitions to energy dense diets [30]. We assessed HFHS-induced changes in EE outcomes as the difference in the 7-day HFHS minus 7-day LFD data. In Figure 6A, 20°C female mice had a ~75% greater HFHS-induced increase in TEE compared to males (p<0.0001), and a 2.8-fold greater increase compared to 30°C females (p<0.0001). Interestingly, no difference was observed between male mice due to temperature. Further, diet-induced changes in the major components of TEE were also observed. Diet-induced REE in 30°C male and female mice was 47% and 69% lower, respectively, compared to 20°C (Figure 6B, p<0.0001). Additionally, 20°C females had ∼25% greater diet-induced REE compared to males (p<0.002). Figure 6D depicts the daily increase in REE due to HFHS feeding and demonstrates the rapid and sustained responses observed across the 7-day intervention. Finally, a main effect of temperature is observed for diet-induced change in NREE in part due to the lack of change in mice at 30°C (Figure 6C, p<0.0001). Furthermore, 20°C female mice demonstrated ∼40% less reduction in NREE due to HFHS feeding compared to males (p<0.0001). These data demonstrate that sex, baseline differences in EE, and diet interact to impact metabolic flexibility and adaptive thermogenic responses in EE in mice.

**Figure 6:**
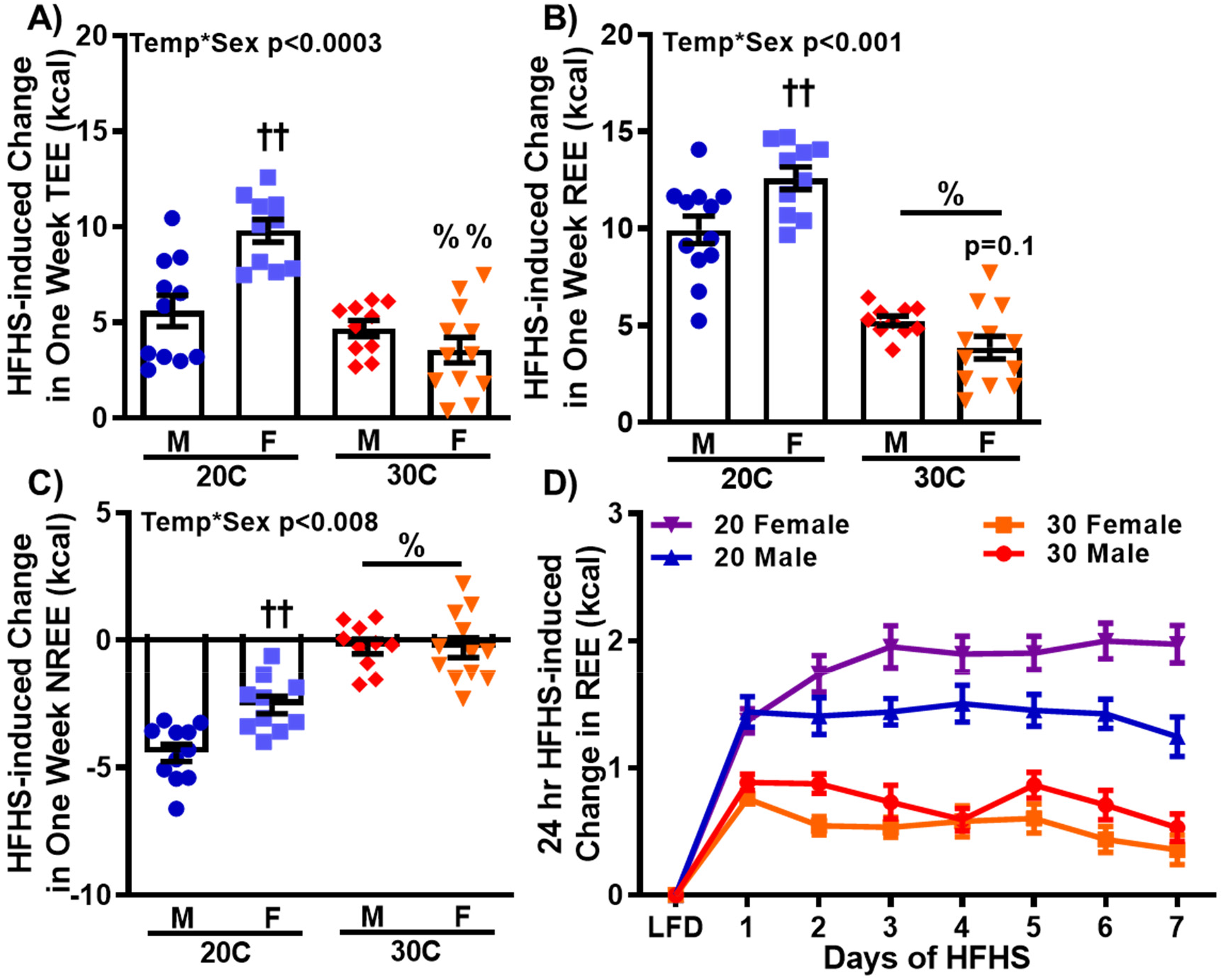
Energy metabolism and sex interact impacting high-fat/high-sucrose-induced changes in energy expenditure. HFHS-induced changes in A) TEE, B) REE, and C) NREE were calculated as the difference between the one week HFHS values and the one week LFD values. D) Daily HFHS-induced change in REE above LFD REE. Values are means ± SEM. n=10-16. % p<0.05 main effect of 20°C vs. 30°C, †† p<0.05 male vs. female within temperature by diet group, %% p<0.05 20°C vs. 30°C within sex by diet group.

### 3.6 Activity Components of Energy Metabolism

Under the sedentary cage conditions of the current study, all activity EE (AEE) would represent non-exercise spontaneous activity. AEE is calculated as the difference in 7-day NREE and the 7-day TEF for each diet (Figure 7A). As TEF is calculated based on food intake and macronutrient composition [27], the data is relatively similar to EI (Figure 1B) with the primary findings being reduced TEF due to temperature for both LFD and HFHS (p<0.0001) and greater for all HFHS groups (p<0.002). HFHS reduced AEE (Figure 7b) regardless of sex (∼40% and ∼15%, 20°C and 30°C, respectively, p<0.0005). However, the reduction in AEE for 30°C males and females was ∼60% greater compared to 20°C mice (p<0.0001). Interestingly, the observed differences in AEE were not associated with any main effect differences in activity level (Figure 7C). This would suggest a difference in the energy cost of movement (CoM) (Figure 7D), which is calculated here as the AEE per meter of movement in the cage. Interestingly, females had lower CoM in all comparisons (p<0.01), suggesting that female mice are inherently more efficient during cage-based movement. Also, HFHS reduced CoM in male (38%) and female (31%) 20°C mice compared to LFD (p<0.0001 & p<0.006, respectively). Male and female 30°C HFHS-fed mice had 30% (p<0.01) & 50% (p<0.005) greater CoM (respectively) compared to 20°C showing that housing temperature impacts energy efficiency of movement robustly. Importantly, while these data highlight sex differences in the EE phenotypes during HFHS, there is no obvious association between activity or AEE with changes in body weight or body composition with acute HFHS feeding.

**Figure 7:**
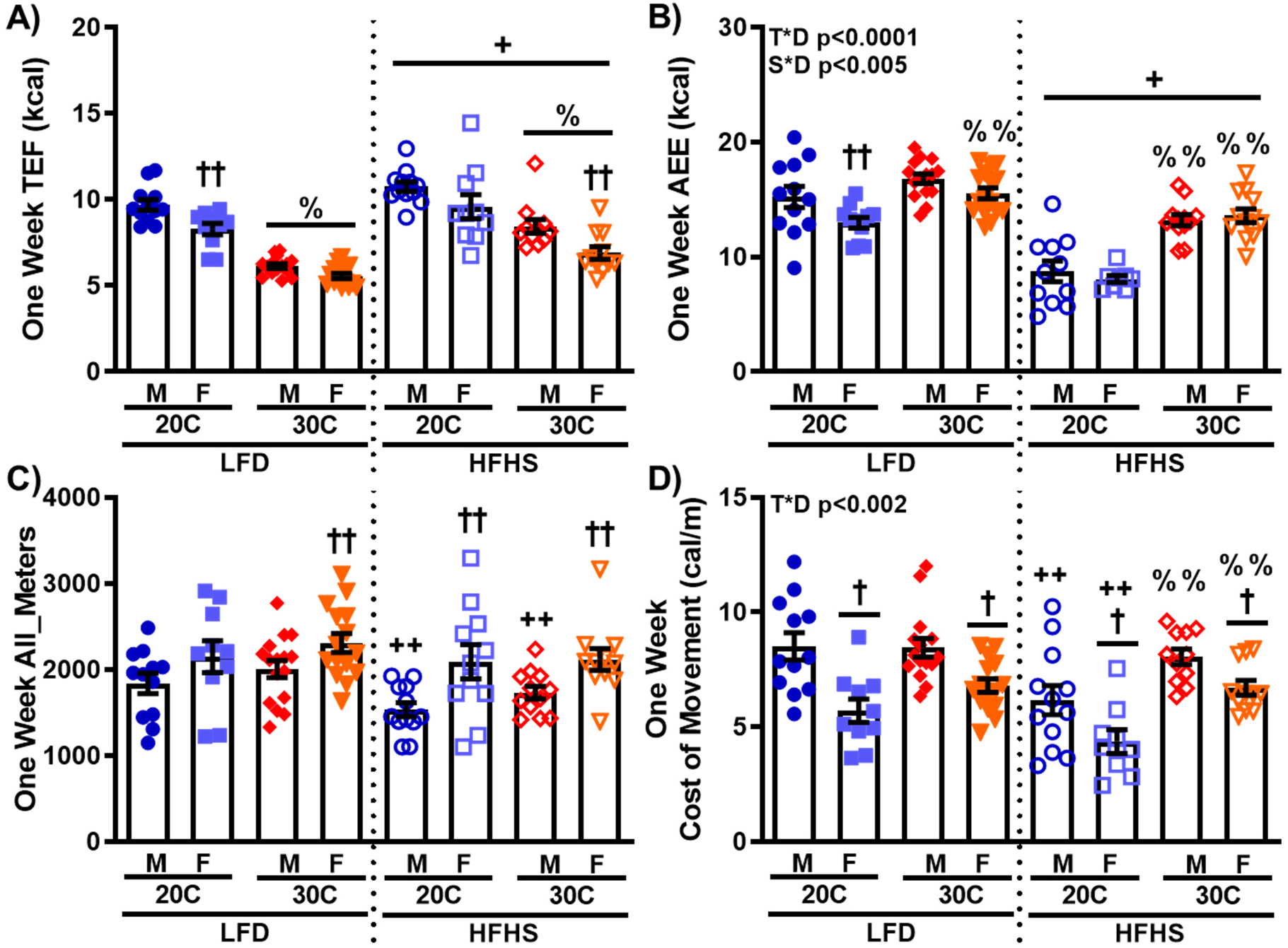
Different ambient housing temperatures does not result in different activity levels, and activity energy expenditure does not associate with the observed changes in body weight or body composition. A) Thermic effect of food (TEF) was determined from weekly food intake and macronutrient composition of each diet, and B) activity energy expenditure (AEE) was calculated as the difference in NREE and TEF (n=9-16). Total cage activity was determined as (C) All_Meters (n=10-16), and the (D) cost of movement (CoM) was calculated as the AEE divided by the cage activity (n=9-15). Values are means ± SEM. % p<0.05 main effect of 20°C vs. 30°C, + p<0.05 main effect of LFD vs. HFHS, † p<0.05 main effect of male vs. female, †† p<0.05 male vs. female within temperature by diet group, %% p<0.05 20°C vs. 30°C within sex by diet group, ++ p<0.05 LFD vs. HFHS within temperature by sex group.

## 4. DISCUSSION

Energy expenditure putatively plays a fundamental role in driving both susceptibility for weight gain and treatment of obesity. However, the direct assessment of the role of EE on EB and weight gain is complicated by potentially confounding factors produced by the common experimental tools available (e.g. – physical activity, chemical uncouplers, etc.). There have also been limited studies examining the links between sex, EE, and weight gain regulation. As a novel experimental tool to assess the independent role of EE on weight gain, we have utilized differing ambient housing temperatures (20 vs. 30°C) to modulate EE. Our primary findings are that housing temperature induced divergence in TEE and REE, and interacted with acute HFHS feeding to produce sexually dimorphic changes in body weight and body composition in C57Bl/6J mice. While male mice with lower EE gained more weight during 7-day HFHS feeding than their counterparts with higher EE, no difference was observed between female mice groups with different EE. Interestingly, all male mice, and female mice with lower EE gained the same relative proportion of weight as fat, however, the female mice with higher EE gained primarily FFM. The changes in adiposity during HFHS feeding were highly associated with the observed EB. Further, the reduced diet-induced adiposity in female mice with greater EE was associated with enhanced HFHS diet-induced changes in RQ and non-shivering thermogenesis. Importantly, while total activity was higher in females, neither activity-induced EE or feeding patterns appear to be associated with sex differences in weight gain and body composition.

EB is not a simple static equation but is rather a dynamic system with EI and EE continually regulating one another to influence thermoregulation and body mass [6–8]. It has been proposed that mammalian physiology has evolved such that optimal maintenance of EB and body weight is achieved at higher levels of EE [6–8]. Jean Mayer was the first to demonstrate in rodents and humans that EB is more highly regulated at higher levels of EE (as increased physical activity), establishing a coupling of EI and EE within certain limits [31, 32]. This work has been extended to describe a zone in which EI and EE are highly coupled (regulated zone), below and above which the components become uncoupled (unregulated zone) [33]. It is proposed that more individuals live within this unregulated zone as a consequence of rising levels of sedentary behavior and physical inactivity. In the current study, greater EE was associated with lower EB during HFHS feeding. Furthermore, the greater weight adjusted EE observed in female mice was associated with reduced EB compared to males housed at both ambient temperatures. In our previous work in rats selectively bred for divergence in intrinsic aerobic capacity, male rats with higher weight adjusted EE also had lower EB following the transition to HFHS diet [34]. In this study, the reduced EB in the 20°C mice was due in part to the smaller HFHS-induced increases in EI compared to 30°C (∼45% vs ∼90%, respectively). While EI during HFHS feeding was lower in female mice at both temperatures, this difference was likely entirely due to the sex difference in body weight. Weight adjusted EI was not different by sex in any of the temperature by diet groups. Changes in food intake patterns or behavior may impact EI and weight gain, however; only the 30°C mice with lower EE had significantly higher food intake on the 7-day HFHS compared LFD (Supplemental Figure 1). Further, while differences in feeding behavior during short-term exposure to the HFHS diet between mice with different EE were observed (Supplemental Figure 1), no association between feeding patterns and the observed HFHS-induced weight gain was observed. These data support recent human findings where greater maintenance of EB exists at higher EE levels due to enhanced EI regulation impacting weight gain [35–37].

Between 60 – 80% of weight gained during most periods of positive EB is fat mass [9]. Considerable sexual dimorphism has been observed in the amount and anatomical location of fat mass gained during hypercaloric conditions [38, 39]. In general, women tend to have higher percent body fat than men, and greater fat deposition in subcutaneous depots. Further, while pre-menopausal women have considerable protection from numerous metabolic disease states [39], the prevalence of overweight/obesity is higher in women in all age groups [40]. Chronic *ad libtum* high-fat diet rodent studies of varying lengths have demonstrated that males and females gain similar amounts of body weight, with females having greater body fat percentages [41, 42]. As expected, we observed less fat mass gain during HFHS feeding in mice with lower EB. Notably, females with higher weight adjusted EE and lower EB gained less fat mass than males. Our previous rat work also showed that higher weight adjusted EE was associated with reduced HFHS induced gains in fat mass [34, 43]. In humans, recent work demonstrated that low levels of physical activity EE are associated with reduced coupling of EI and increased fat mass gain [44]. Interestingly, 20°C female mice with increased EE were the only group to gain fat-free mass rather than fat mass during HFHS feeding. The observed increase in fat-free mass in the 20°C females represented over 80% of the body weight gain, compared to only ∼30% for the other three groups. Relatively few studies have focused on changes in fat-free mass during overfeeding studies, but it is generally attributed to changes in total body water (reviewed in [45]). Though not reported, lean mass was determined during the body composition analysis and showed the same outcomes as calculated fat-free mass. Importantly, differences in body composition driven by ambient temperature and sex were only apparent during the HFHS feeding, illustrating the importance of EE to limit excessive weight gain and adiposity during consumption of energy dense diets [6–8].

The capacity to adapt energy metabolism to diet macronutrient composition through changes in substrate utilization and non-shivering thermogenesis likely drive susceptibility for obesity and metabolic disease and are thus a focus of treatment [10, 29, 30, 46–49]. Metabolic flexibility was initially described as the capacity to alter fuel utilization during the transition from fasting to fed states [50]. The initial investigation demonstrated that skeletal muscle of fasted lean subjects utilized more fat (lower RQ) and responded to insulin infusion with rapid increases in glucose utilization (higher RQ, metabolically flexible), while the obese subjects utilized less fat during fasting and did not increase glucose utilization in response to insulin (metabolically inflexible). In this study we observed that male and female mice with lower EE due to 30°C ambient housing have dramatically reduced metabolic flexibility on LFD compared to 20°C mice, and virtually no within day metabolic flexibility during the HFHS feeding. The LFD findings are significant in that no difference in average 24 hr RQ was observed between the temperature groups; however, the substantial difference in metabolic flexibility is driven by higher RQ in the dark cycle and lower RQ in the light cycle of the 20°C mice (data not shown). Based on differences in HFHS-induced weight gain between the 20°C and 30°C mice, our data suggests that reduced within-day metabolic flexibility is a predisposing factor to weight gain. Our data also shows that the degree of metabolic flexibility is highly associated with EE. This concept is supported by similar work demonstrating differences in metabolic flexibility in two mice strains with different susceptibility to obesity [51]. The definition of metabolic flexibility has expanded to encompass adaptation of substrate utilization in response to changes in dietary macronutrient composition, including acute high-fat diet feeding [29]. Our previous work showed that rats with increased aerobic capacity and greater weight adjusted EE have a greater change in fat utilization (lower RQ) and greater total fatty acid oxidation during HFHS feeding [34]. In this study, while the HFHS feeding resulted in an immediate reduction in average daily RQ in all mice, the increased EE of 20°C mice was associated with greater change in average daily RQ following the dietary transition. Recently, the arcuate nucleus and ventromedial nucleus of the hypothalamus have been identified in the central control of metabolic flexibility during high-fat diet [52, 53], and regulation associated changes in EE [54]. Additional work is necessary to decipher how substrate availability and utilization interact with EE to alter central regulation of energy homeostasis and weight gain.

Non-shivering adaptive thermogenesis is the increased production of heat in response to either cold or diet stimuli [48]. Since the observation of thermogenic adipose depots in humans, and the discovery of skeletal muscle thermogenic capacity independent of contraction, many laboratories have explored the mechanisms of central regulation and activation required to produce heat uncoupled from oxidative phosphorylation as a fulcrum for obesity treatment (reviewed in [30, 48, 49]). While the efficacy of these approaches is still in question, understanding how diet-induced increases in EE may limit weight gain during acute energy surfeit is important [55]. Rodent work has highlighted the potential importance of diet-induced non-shivering thermogenesis through numerous findings of increased susceptibility or protection for diet-induced weight gain following knockout [56–59] or overexpression [60] of genes involved in various thermogenic pathways. In human subjects the major determinant for individual diet-nduced thermogenesis is energy content [61]. However, obese subjects have lower diet-induced thermogenesis compared to lean [62] implicating reduced adaptability in EE as a mediator of weight gain. In this study we observed that 20°C females with higher EE having greater 1-week HFHS diet-induced thermogenesis than 20°C males or the 30°C females. This greater adaptation of EE in the females to the HFHS diet was apparent in both the TEE and REE. In contrast, only the REE component adapted differently in males housed at 20°C and 30°C. This was due, in part, to a greater diet-induced reduction in NREE in 20°C males than both 20°C females and 30°C males. The HFHS-induced non-shivering thermogenesis observations reported here are the first to show that EE and sex interact to alter systemic thermogenic response to energy dense diet. Estrogen signaling in the hypothalamus is important for regulation of energy homeostasis in females (Reviewed in [63, 64]), particularly in the ventromedial hypothalamus which is involved in regulation of diet-induced non-shivering thermogenesis. However, further studies are necessary to determine if increased diet-induced non-shivering thermogenesis in female mice is obligatory for their phenotype of reduced weight gain, and if estrogen signaling is critical for both processes. Overall, these findings support a role for diet-induced non-shivering thermogenesis in the reduced diet-induced fat mass gain in 20°C mice with increased EE.

### 4.1 Limitations

Despite the wide breadth of energy metabolism data collected during these experiments, several potentially confounding factors and limitations should be considered. First, while the C57Bl/6J mouse strain is extensively utilized in obesity studies, the use of other inbred and outbred mouse strains for future studies, particularly related to assessment of sex differences, are necessary. Second, the increased EE of mice at sub-thermoneutral ambient temperatures is primarily mediated by centrally regulated non-shivering thermogenic pathways in adipose and skeletal muscle. These pathways differ from those potentially activated through increased physical activity or exercise and may confound the findings. Further, from the data herein we can not determine the magnitude of activation of the different non-shivering thermogenic tissues, which could potentially differ by baseline EE or sex. Third, the assessment of diet-induced weight gain in 9 – 11 week old mice could be confounded by the previously observed dependence of weight gain on age of diet initiation and sex [41]. Fourth, previous mouse work has demonstrated that male mice defend different body core temperatures at different ambient temperatures [27, 65]. The lack of these thermal biology data prevents a comprehensive dissection of sexual differences in energy metabolism, and the impact on metabolic responses to short-term HFHS feeding. Finally, the lack of fecal energy excretion data prevents the calculation of net energy intake during both the LFD and HFHS feeding, and potentially confounds the calculation of EB.

### 4.2 Conclusions

Because the prevention of weight gain is putatively easier than weight loss [6], it is critical that mechanisms underlying the protection or susceptibility to episodic weight gain during hypercaloric conditions be elucidated. This study used ambient temperature to determine if higher or lower EE in male and female mice would change metabolic adaptations and weight gain during a transition to acute HFHS feeding. Here we demonstrate that baseline EE and sex interact to impact diet-induced changes in body composition and weight gain. This interaction is a result of the observed inverse relationship between fat mass gain and weight-adjusted TEE, as well as, diet-induced non-shivering thermogenesis. These data offer further support that at higher levels of EE, there is enhanced coupling of EI to EE during HFHS, resulting in reduced positive energy balance and reduced gains in weigh and adiposity. Additionally, these data demonstrate that EE level plays a role in the composition of weight gained by female mice during acute HFHS feeding. Finally, these findings have increased significance when one considers that the vast majority of obesity research conducted in mice occurs at sub-thermoneutral housing, near, the 20°C temperature.

## Author Contributions

Author contributions: EMM, JPT, conception and design of research; EMM, RDN, JAA, CSM performed experiments; EMM, QX, DCK, RPS, JRBL, analyzed data; EMM, RDN, JAA, CSM, RPS, JRBL, JAC, JPT, interpreted results of experiments; EMM prepared manuscript; EMM, RDN, JAA, CSM, QX, DCK, RPS, JRBL, JAC, JPT edited and revised manuscript.

