## Supplemental Table and Figure for "Divergent Energy Expenditure Impacts Mouse Metabolic Adaptation to Acute High-Fat/High-Sucrose Diet Producing Sexually Dimorphic Weight Gain Patterns"

**Supplemental Table A.1. Anthropometrics**

| | 20 | | 30 | |
|---|---|---|---|---|
| | **Male** | **Female** | **Male** | **Female** |
| *Body Weight* | | | | |
| **Initial** | 24.13 ± 0.40 | 18.35 ± 0.36$^{†}$ | 23.93 ± 0.28 | 18.33 ± 0.28$^{†}$ |
| **Day 7 – End of LFD** | 24.21 ± 0.34 | 18.67 ± 0.31$^{†}$ | 25.26 ± 0.35$^{\%}$ | 19.46 ± 0.24$^{†,\%}$ |
| **Day 14 – End of HFHS** | 26.91 ± 0.52 | 21.13 ± 0.48$^{†}$ | 28.98 ± 0.51$^{\%}$ | 21.76 ± 0.40$^{†,\%}$ |
| *Body Composition* | | | | |
| **Fat Mass (g)** | | | | |
| - **Initial** | 1.83 ± 0.11 | 1.88 ± 0.15 | 2.14 ± 0.10 | 1.71 ± 0.10$^{††}$ |
| - **Day 7** | 1.74 ± 0.17 | 1.53 ± 0.08$^{†}$ | 2.47 ± 0.14$^{\%}$ | 1.84 ± 0.09$^{†,\%}$ |
| - **Day 14** | 2.97 ± 0.36 | 1.45 ± 0.26$^{†}$ | 5.18 ± 0.30$^{\%}$ | 3.28 ± 0.28$^{†,\%}$ |
| **Fat-free Mass (g)** | | | | |
| - **Initial** | 22.30 ± 0.41 | 16.25 ± 0.30$^{†}$ | 21.91 ± 0.24 | 16.62 ± 0.31$^{†}$ |
| - **Day 7** | 22.72 ± 0.43 | 17.21 ± 0.33$^{†}$ | 22.79 ± 0.26 | 17.63 ± 0.24$^{†}$ |
| - **Day 14** | 23.54 ± 0.34 | 19.01 ± 0.38$^{†}$ | 23.80 ± 0.27 | 18.48 ± 0.20$^{†}$ |
| **Body Fat Percent** | | | | |
| - **Initial** | 7.60 ± 0.45 | 10.35 ± 0.83$^{††}$ | 8.88 ± 0.0.37 | 9.38 ± 0.59 |
| - **Day 7** | 7.13 ± 0.68 | 8.15 ± 0.45 | 9.71 ± 0.45$^{\%}$ | 9.44 ± 0.48$^{\%}$ |
| - **Day 14** | 12.37 ± 0.99 | 9.99 ± 0.71$^{†}$ | 17.75 ± 0.78$^{\%}$ | 14.91 ± 1.08$^{†,\%}$ |

All values expressed as mean ± SEM (n=11-16). % $p<0.05$ main effect of 20ºC vs. 30ºC, † $p<0.05$ main effect of male vs. female, †† $p<0.05$ male vs. female within temperature by diet group.

**Supplemental Figure 1**

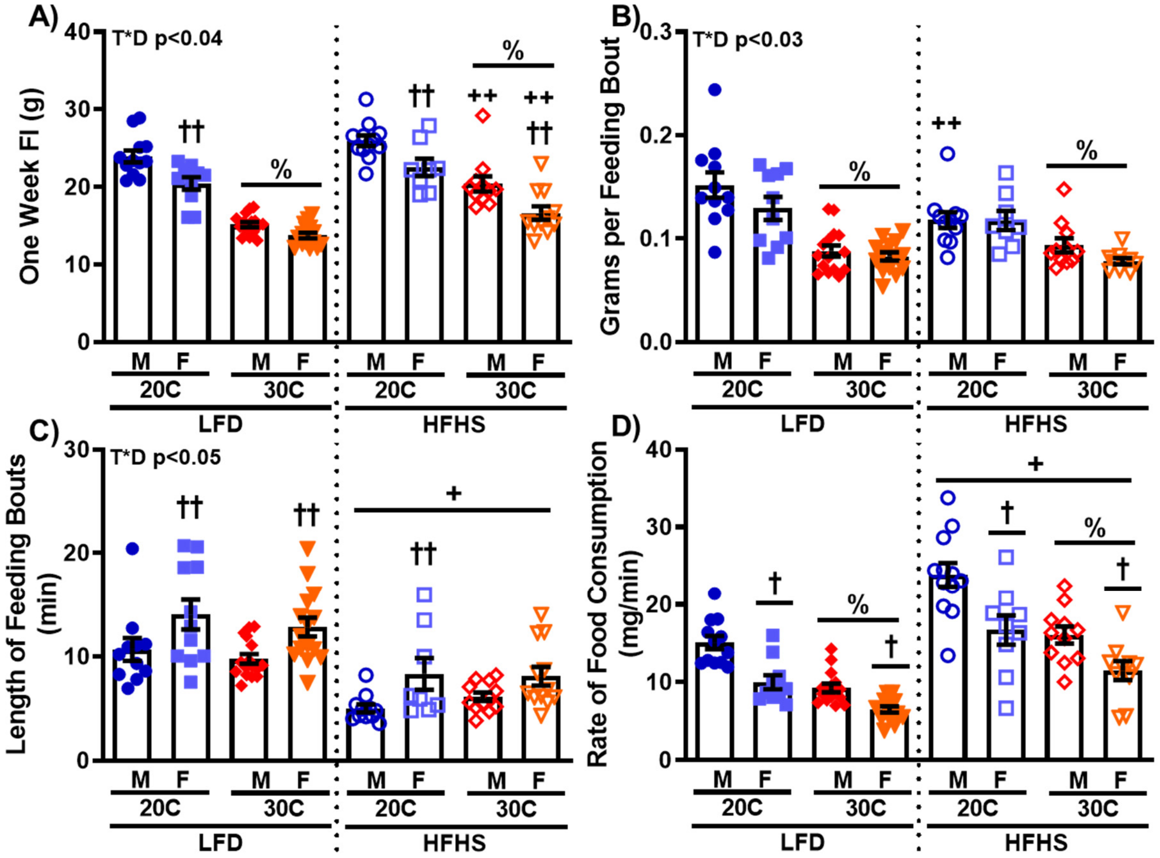


**Supplemental Figure 1: No apparent association exists between feeding patterns and the observed changes in body weight or body composition.** A) One week food intake (FI). Feeding patterns were assessed as: B) gram per feeding bout, C) length of feeding bouts, and D) rate of food consumption. Values are means ± SEM. n=8-16. % p<0.05 main effect of 20ºC vs. 30ºC, + p<0.05 main effect of LFD vs. HFHS, † p<0.05 main effect of male vs. female, †† p<0.05 male vs. female within temperature by diet group, ++ p<0.05 LFD vs. HFHS within temperature by sex group.
